# Host inducible-HSP70A1A is an irresistible drug target to combat SARS-CoV2 infection and pathogenesis

**DOI:** 10.1101/2023.05.05.539661

**Authors:** Prerna Joshi, Swati Garg, Shailendra Mani, Kamini Jakhar, Manisha Marothia, Rumaisha Shoaib, Shreeja Biswas, Jhalak Singhal, Ankita Behl, Amandeep Kaur Kahlon, Maxim Shevtsov, Pramod Garg, Shailja Singh, Anand Ranganathan

## Abstract

One of the fundamental mechanisms developed by the host to contain the highly infectious and rapidly proliferating SARS coronavirus is elevation of body temperature, a natural fallout of which is Heat Shock Protein (HSP) over-expression. Here, for the first time, we demonstrate that the SARS-CoV-2 virus exploits the host Hsp70 chaperone for its entry and propagation and blocking it can combat the infection. SARS-CoV-2 infection as well as febrile temperature enhanced Hsp70 overexpression in host Vero E6 cells. In turn, Hsp70 overexpression elevated the host cell autophagic response that is a prerequisite for viral propagation. Suppressive and prophylactic treatment of Vero E6 cells with HSP70 inhibitor PES-Cl, a small molecule derivative of Pifithrin μ, abrogated viral infection more potently than the currently used drug Remdesivir by suppressing host HSP70 and autophagic response. In conclusion, our study not only provides a fundamental insight into the role of host Hsp70 in SARS-CoV-2 pathogenesis, it paves the way for the development of potent and irresistible anti-viral therapeutics.

## Introduction

The COVID-19 pandemic, caused by SARS-CoV-2 coronavirus, has inflicted a serious and debilitating toll on world health and economy. Remdesivir, the only drug widely used against SARS-CoV-2 during the pandemic did not show promising results advocating development of new therapeutics. Like all other RNA viruses, coronaviruses are reliant on host cellular pathways for their propagation [1]. Infection is characterized by symptoms like high fever, cough, lung injury, metabolic acidosis [2] and multiple organ dysfunction [3], depending upon viral load and severity of the disease. Infection mediated high fever results in the overexpression of genetically conserved numerous stress proteins including the Heat shock protein 70 (Hsp70 or Hsp70A1A) chaperone. Hsp70 is specifically known for its pro-viral effects in Zika [4], Rabies [5], Influenza A [6], and Dengue [7] viral infections, so much so that JG40, JG18 and VER155008 allosteric inhibitors of Hsp70 [7] have been shown to block Dengue virus infection in human Huh7 cells. Herein, we investigated the role of host Hsp70 in SARS-CoV-2 pathogenesis, in the belief that SARS-CoV-2, too, has evolved survival strategies by sequestering cellular pathways like heat shock response and autophagy and alterations in cellular microenvironment induced by Hsp70 expression may facilitate viral infection.

## Results

We examined the expression levels of host Hsp70 in SARS-CoV-2 (Wuhan isolate NC_045512) infected Vero E6 cells (Kidney epithelial cells of *Cerocopithecus aethiops*) by Immunofluorescence Assay (IFA) using HSP70A1A-specific monoclonal antibody [8]. Increased fluorescence intensity of Hsp70 in IFA indicate that SARS-CoV-2 infection elevates the levels of host Hsp70 in Vero E6 cells (Figure 1A). We then assessed the Hsp70 expression, by IFA and Western blot analysis, in febrile conditions (40°C) and mimic viral infection. Induction of higher Hsp70 expression levels was observed by IFA in Vero E6 cells following heat shock similar to virus infected cells (Figure 1B (i)). Hsp70 was uniformly localised throughout the cell, but at some places, high intensity was observed on the surface of cell membrane. We also analysed Hsp70 expression in Vero E6 cells at different time points (1,3,6,12 and 24h) after heat shock using Western blotting. Data suggests that Hsp70 expression starts increasing 6h after heat shock and increases up to 30-fold in 24 hours post heat shock (Figure 1.B (ii)). Next, we analysed the role of Hsp70 in modulating cellular pathways like Autophagy which is required for SARS-CoV2 replication within host cells [9]. Autophagy markers such as, (i) Beclin 1, that binds ATG’s and initiate phagophore formation;(ii) Atg5, that forms complex with Atg12 and (iii) LC3 which yields LC3-II upon modification, were assessed by Western blot analysis in heat shock induced Hsp70 overexpressing Vero E6 cells. Starvation induced autophagy, by incubation in Hank’s buffered salt solution (HBSS), was used as positive control during the experiment. As shown in Figure 1C(i) heat shock induced ∼6-fold Beclin-1 expression while LC3 increased upto 2-fold as compared to control Vero E6 cells. To further validate the connection between Hsp70 expression and autophagy initiation, we treated Vero E6 cells with 15μM of PES-Cl, a known inhibitor of the Substrate-binding domain (SBD) of Hsp70 [10], 1h after heat shock and analysed the autophagy markers. Treatment with PES-Cl have no cytotoxic effect on Vero E6 cells but significantly reduced the levels of autophagy markers (Figure 1C (ii, iii), Supplementary figure 1, 2). This suggests that Hsp70 expression regulates initiation of autophagy in Vero E6 cells. Hsp70 expression on cell membrane suggests the requirement of chaperonin activity near surface. SARS-CoV-2 viral entry is mediated by interaction between receptor binding domain (RBD) of spike protein and human Angiotensin converting enzyme receptor (ACE-2) at cell membrane [12]. Thus, we assessed Hsp70 interaction with ACE-2 and RBD using Surface Plasmon Resonance (SPR) and in silico docking (Figure 1D(i), Supplementary figure 3, 4, 5, 6, Supplementary table 1). As already known, ACE-2 and RBD strongly interact with each other, hence we observed K_d_ value of 4.63μM. The K_d_ value for ACE-2-Hsp70 binding was 7.04 μM suggesting strong interaction between human ACE-2 and Hsp70. While Hsp70 interaction was not observed with RBD protein of SARS-CoV2 (Figure 1D(i)). This suggest that Hsp70 may play chaperonin role near the cell membrane and help proper folding of human ACE-2 receptor to facilitate viral entry. To investigate the potential role of Hsp70 in SARS-CoV2 infection, we treated SARS-CoV-2 infected Vero E6 cells with different concentration of PES-Cl. Viral load was assessed in culture supernatants by qRT-PCR. A 15μM concentration of PES-Cl approximately inhibited 100% viral load (Figure 1E). Remdesivir, a known antiviral, showed about 70% inhibition at same concentration suggesting much more efficacy of PES-Cl in reducing viral load. Since Hsp70 interacted with ACE-2, we inquired its role in virus entry. Vero E6 cells were pre-treated with PES-Cl, Remdesivir and mAb-HSP70, washed and incubated with virus. PES-Cl demonstrated highly reduced viral load in culture supernatants as compared to Remdesivir. The mAb-HSP70 does not show any inhibition suggesting that Hsp70 is not involved in any physical interaction rather its chaperonin activity is involved in viral infection. Overall, our data demonstrates inhibition of Hsp70 by PES-Cl inhibits virus invasion and reduces production of infectious SARS-CoV2 particles. Collectively, our results suggest that Hsp70 is a crucial player that regulates cellular processes like host autophagy during SARS-CoV-2 infection and is an excellent target for anti-viral therapeutics development (Figure 1F).

**Figure 1:**
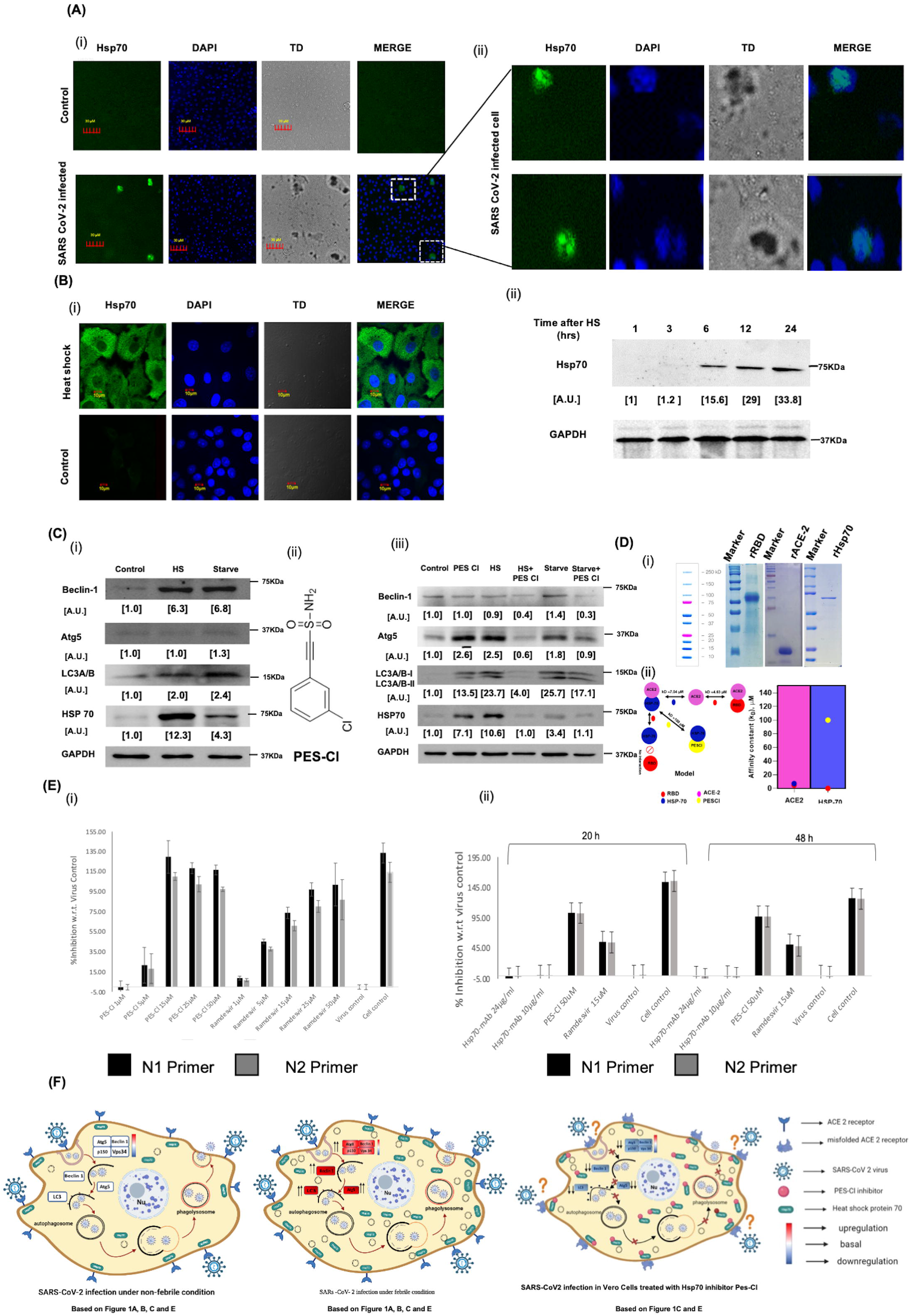
(A) (i) Intracellular expression of Hsp70 in Vero E6 cells following 48h viral infection. Cells were stained for Hsp70 (green) and counterstained stained with DAPI (blue). (ii) Zoomed in images of highlighted cells in figure (i). Scale bar 30μm. (B) Induction of temperature dependent Hsp70 expression in Vero E6 cells. (i)Upper panel shows confocal imaging of heat shock induced Vero E6 cells. Heat shock was given at 40°C for 1h. Cells were stained for Hsp70 (green) and cell nuclei stained with DAPI (blue). Lower panel shows control cells without heat shock. Scale bar 10μm. (ii) Time dependent Hsp70 expression in Vero E6 cells was assayed by Western blotting at indicated timepoints following heat shock. Hsp70 expression was quantified by ImageJ software and GAPDH was assayed as a housekeeping control. (C) Hsp70 expression regulates the initiation of autophagy. (i)Upon heat shock (HS) at 42oC, expression of autophagy markers beclin-1, ATG5 and LC3 A/B increased as compared to control cells (no heat shock). Starvation (6h in HBSS) induced Vero E6 cells were used as Positive control. (B)Treatment with PES-Cl, an inhibitor of Hsp70 significantly reduced the expression of autophagy markers. GAPDH was assayed as a housekeeping control. (D) Surface Plasmon resonance reveals positive interaction between human Hsp70 and ACE-2 receptor. (i) Coomassie stained SDS-PAGE analysis of the Ni-NTA purified His-tagged proteins (RBD of spike protein, ACE-2 and Hsp70) M: marker, 1: Purified His-tagged RBD (26.3KDa), 2: Purified His-tagged ACE-2(13.1KDa), 3: Purified His-tagged Hsp70 (70KDa). (ii)Simplified model depicting the interaction between Hsp70 and ACE-2 while negative interaction between receptor binding domain of spike protein (RBD) and Hsp70. Graph depicting SPR based interaction analysis between ACE-2-RBD; ACE-2-Hsp70 and Hsp70-PES-Cl. (E) Hsp70 inhibitor PES-Cl inhibits infectious SARS-CoV2 production. (i) Vero E6 cells were infected with SARS-CoV2, washed, and replenished with medium containing indicated drugs. (ii) To assess role of Hsp70 in SARS-CoV2 entry, Vero E6 cells were pretreated with indicated drugs followed by virus infection. Viral load was assessed in culture supernatants after 48h. Remdesivir was used as a positive control. (F) Proposed model for the role of Hsp70 in autophagy mediated SARS-CoV2 infection. Under normal temperature conditions, HSP70 regulates basal autophagy and helps viral replication. Febrile conditions or infection induces Hsp70 expression leading to enhanced autophagy with increased viral load. Inhibiting the Hsp70 by PES-Cl downregulates autophagy pathway and hence viral infection.

## Discussion

Efficient counter-measures to tackle SARS-CoV-2 are still being developed, with identification of host proteins that facilitate viral infection increasingly being recognized as a promising new antiviral strategy. That said, traditional antivirals targeting viral proteins leads to rapid mutations in the latter rendering these therapies ineffective. Therefore, targeting host proteins that are essential for viral pathogenesis would avoid the spectra of resistance and help develop broad-spectrum antivirals.

Under normal temperature conditions, Hsp70 regulates basal autophagy and helps viral replication. Febrile conditions due to heat shock or viral infection induces Hsp70 expression. Elevated Hsp70 levels inside the cell further upregulate autophagy markers like Beclin-1 and LC3 suggesting initiation of autophagosome formation. Treatment with PES-Cl showed significant decline in levels of early autophagy markers and SARS-CoV-2 infection with a potency much greater than the widely used antiviral drug Remdesivir. Moreover, Remdesivir targets the viral protein hence can be modified by virus leading to emergence of resistance. But, targeting highly conserved host protein like Hsp70 for antiviral development is therefore an irresistible strategy.

Autophagosomes can provide a platform for viral replication congregating replication machinery and further prevent detection of viral RNA by the innate immune system. Furthermore, studies have shown regulation of autophagy by Hsp70. Our data is consistent with previous studies that elucidated active involvement of autophagy in SARS-CoV2 infection [13] [14] [9]. We hypothesize that this is a survival strategy of the virus wherein it hijacks cellular defense machinery for its own replication and protects itself from the host innate immune system. SARS-CoV-2 has thus evolved survival strategies to fine-tune cellular pathways like heat shock response and autophagy so that it can harness benefits from the host without getting destroyed. Targeting these mechanisms to develop therapeutics against coronavirus disease is therefore a promising way forward. Pandemics like COVID-19 demands treatments that are both preventive and suppressive. Thus, targeting HSP70 by PES-Cl provide a novel therapeutic strategy to restrict viral entry and replication.

## Acknowledgements

We would like to dedicate this manuscript to colleagues and patients worldwide who suffered and succumbed to SARS-CoV2 infection. We thank Central Instrumentation Facility (CIF) of Special Centre for Molecular Medicine, JNU. We also thank Advanced Instrumentation and Research Facility (AIRF), JNU, New Delhi for confocal microscopy. This work is supported by funding from Department of Science and Technology, SERB IRHPA IPA/2020/000007 (AR, SS) and Indian Council of Medical Research, NER/84/2022-EDC-I (SS). Funding from the National Bioscience Award from DBT to SS is acknowledged. The graphical model was designed with BioRender (BioRender.com). This manuscript has been screened by the anti-plagiarism software.

