## Supplementary information for "Host inducible-HSP70A1A is an irresistible drug target to combat SARS-CoV2 infection and pathogenesis"

### Material and Methods

**Cell culture:** VeroE6 (Kidney epithelial cells of Cerocopithecus aethiops) obtained from National Center for Cell Sciences (NCCS) were maintained in Dulbecco's Modified Eagle Medium (DMEM)(HIMEDIA) supplemented with 10% heat-inactivated fetal bovine serum (FBS,Gibco ThermoFischer Scientific) ,1% Penicillin-Streptomycin (Gibco ThermoFischer Scientific, Waltham, MA, USA) and 1% MEM Non-Essential Amino Acids Solution (100X) (Gibco ThermoFischer Scientific, Waltham, MA, USA). Cells were cultured at 37°C with 5% CO2.

**Virus stock and SARS-CoV-2 infection**: SARS-CoV-2 was obtained from Biodefense and Emerging Infections Research Resources Repository (BEI resources, NR52281), amplified in Vero E6 cells in the biosafety level 3 (BSL3) facility of THSTI, India, titrated, and stored frozen in aliquots. For SARS-CoV-2 infection studies, the stock virus was diluted in serum-free medium to 200ul volume consisting of 400 50% tissue culture infective doses ($TCID_{50}$) and added to a Vero E6 cell monolayer (seeded in a 24-well plate) for 1 h at 37°C, supplemented with 5% $CO_2$. At 1 h post-incubation, the infection medium was removed, cells were washed twice with 500 µl of serum-free medium, and fresh DMEM plus 2% FBS (maintenance medium) was added. In the case of PES-Cl (Sigma-aldrich) pre-treatment, PES-Cl was added to the culture medium 8h before infection. 8h post treatment with PES-Cl, cells were washed with serum-free medium and SARS-CoV-2 infection was given in Vero E6 cells for 1hr. Following infection, cells were again washed with serum-free medium and maintenance medium was added, and maintained for 20h (in pre-treatment study) and 48 h (in both pre and post-treatment studies) , followed by collection of culture medium for RT-qPCR assay and cells for subsequent experiments .In case of remdesivir, it was added to the culture medium during infection, again added to the complete medium after removal of the infection medium, and maintained for 20h (in pre-treatment study) and 48 h (in both pre and post-treatment studies), followed by collection of culture medium and cells for subsequent experiments.

**RNA Isolation and RT-qPCR assay**: Intracellular RNA was isolated using TRIZOL reagent (MRC, MA, USA), followed by reverse transcription (RT) using a FIREScript cDNA synthesis kit (Solis Biodyne, Estonia). RNA from culture medium was isolated using a Qiagen viral RNA minikit (Qiagen, Germany), followed by RT using a FIREScript cDNA synthesis kit (Solis Biodyne, Estonia). Random hexamers were used in cDNA synthesis. SYBR green-based quantitative real-time PCR (RT-qPCR) was done. Two sets of primers (N1 and N2) of the Nucelocapsid gene were used (as given below) and TaqMan-based RT-qPCR was done by following the protocol suggested by the CDC, USA. Absolute quantification of each gene was carried out to determine viral RNA copy no.s/ml. For this, synthetic single-stranded RNA with known copy number (Merck, EURM019) was used as a reference to generate the standard curve, with each run.

| | | |
|---|---|---|
| **N1- Forward Primer** | | GACCCCAAAATCAGCGAAAT |
| **N1-Reverse Primer** | | TCTGGTTACTGCCAGTTGAATCTG |
| **N1-Probe** | FAM | ACCCCGCATTACGTTTGGTGGACC |
| **N2- Forward Primer** | | TTACAAACATTGGCCGCAAA |
| **N2- Reverse Primer** | | GCGCGACATTCCGAAGAA |
| **N2-Probe** | JOE | ACAATTTGCCCCCAGCGCTTCAG |

**Antibodies and chemical reagents:** Primary antibodies used in Western blotting were Beclin-1 (CST, Danvers, MA, USA; #3495T, 1:1,000), Atg5 (CST #12994T, 1:1,000), LC3A/B (CST #4108S, 1:1,000) and GAPDH (Invitrogen,1:5000). Goat anti-rabbit HRP-conjugated secondary antibody (1:2,500) was purchased from ABclonal (Woburn, MA, USA). Hsp70-specific antibody (cmHsp70.1) was used for western blotting and immunofluorescence analysis. Goat anti-mouse HRP-conjugated secondary antibody was purchased from Invitrogen. Alexa Fluor 488 goat anti-mouse IgG was purchased from Invitrogen. Coverslips were mounted on the slide for immunofluorescence imaging by mounting medium, fluoroshield with DAPI (Sigma).

**Chemical compounds and Antiviral screening assay:** A 10mM stock of Remdesivir (R&D systems) was dissolved in DMSO and stored at -80˚C in aliquots for single use. It was used as a positive control during antiviral assays. PES-Cl (Sigma-Aldrich) was also dissolved in DMSO to make a final concentration of 1mM concentration, aliquoted and stored at -20˚C.Vero E6 cells were seeded in 48 -well plate at a seeding density of $0.05 \times 10^6$ cells/well and grown overnight at 37˚C.Serial dilutions of the compound and Remdesvir (from 50uM to 1uM) were prepared in DMEM and 500ul of each drug (consisting of the desired molarity) was added 8 hours prior to infection. Subsequently, wells were infected with SARS-CoV-2 Wuhan strain in order to evaluate inhibition of infection. Each compound concentration was tested in triplicates and assay plate contained following controls: cell control (mock infection) and virus control.

**Cell Viability assay:** To assess the effect of PES-Cl on VeroE6 cell viability, MTT assay was done. MTT dye [3-(4,5-dimethyl-2-thiazoyl)-2,5-diphenyltetrazolium bromide] (Sigma-Aldrich, USA) was used at a concentration of 5mg/ml. Cells were seeded in a 96-well plate at seeding density of 10,000 cells/100µl.Cells were exposed to varying concentrations of PES-Cl. Cells treated with only media served as a control group. After 48 hours, MTT assay was carried according to manufacturer’s protocol. Briefly, media was removed from inhibitor treated and control Vero E6 cells and cells washed with PBS twice. 20µl of MTT and 100 µl of fresh medium were introduced. Cells were incubated with MTT dye for 2h at 37˚C and then DMSO was added. Absorbance was measured by microplate reader after 30 minutes at 570nm and cell viability percentage calculated. Each experiment was done in triplicates and repeated twice.

**Heat Shock assay**: Vero E6 cells were seeded in 6 well plate at a seeding density of $0.3 \times 10^6$ cells/well. Confluent cell monolayers were then exposed to heat shock for 1 h at 40˚C followed by 37˚C.Cell lysates were collected 1h,3h,6h,12h and 24h post heat shock. Cell lysates were prepared in radioimmunoprecipitation assay (RIPA) lysis buffer (G-Biosciences, St. Louis, MO, USA) and the addition of protease inhibitor cocktail (Roche, Basel, Switzerland).

**Starvation assay**: Confluent Vero E6 cell monolayers were grown in 6 well plate maintained at 37˚C and 5% CO2 overnight. Next day media was removed and cell were washed twice with 1X PBS. Media was replaced with Hank’s buffered salt solution (HBSS)(Cytiva) for 6h to induce starvation and then recovered. Lysates were collected in RIPA lysis buffer after 24 hours.

**Western Blot analysis**: Cells were washed with PBS, trypsinized and pelleted by centrifugation. Cell pellet was lysed in radioimmunoprecipitation assay (RIPA) lysis buffer (G-Biosciences, St. Louis, MO, USA) and lysates were collected. Lysates were centrifuged at 13,000 rpm for 10 minutes at 4˚C.Supernatants were resuspended in 5X SDS loading dye. Equal volumes of lysates were loaded on sodium dodecyl sulfate–polyacrylamide gel electrophoresis (SDS-PAGE), after which proteins were transferred onto the nitrocellulose immunoblot membranes (Bio-Rad, Hercules, CA, USA). The membranes were blocked in 1× PBST buffer containing 0.05% Tween 20 (Sigma) and 5%BSA overnight at 4°CNext day, membranes were washed thrice with 1XPBST, and further incubated with primary antibodies for 2 h at room temperature following three washes (1X PBST) and incubation with HRP-conjugated secondary antibody for another 1 h at room temperature. After three washes, blots were visualized with enhanced chemiluminescence (ECL) kit (Bio-Rad) on the ChemiDoc Imaging System (Bio-Rad). Band intensities were quantified using ImageJ software (NIH, Bethesda, MD, USA).

**Hsp70 Immunolocalization**: Immunofluorescence imaging was performed to evaluate the localization of Hsp70 in Vero E6 cells. Cells were seeded at a density of 0.3 $x10^6$ cells/ml in a 6 well plate with coverslips in each well. Heat shock was given to cells and cells were recovered back at 37˚C.After 24 h cells were washed twice with PBS and fixed with chilled methanol for 30 minutes at room temperature. Cells were then washed with PBS and permeabilized in permeabilization buffer (0.05% Triton,3% BSA in PBS) for 15 minutes followed by Blocking with 3% BSA for 30 minutes. Cells were then incubated with primary antibody (Hsp70 monoclonal antibody) for 45 minutes at room temperature. Following washes with PBST, cells were incubated with Alexa Fluor 488 goat anti-mouse IgG (Invitrogen). Cells were then washed, and coverslips were mounted on the slides with a mounting medium, fluoroshield with DAPI (Sigma). Image was acquired under confocal microscopy. Confocal imaging was performed with Olympus Fluoview FV1000 with 60× objective magnifications.

**Cloning, expression and protein purification of Receptor binding domain and human Angiotensin converting enzyme-2**: For the cloning of Receptor binding domain (RBD) and Angiotensin converting enzyme-2 (ACE-2) in pMTSAra vector, the inserts were prepared through restriction digestion using SnaBI, followed by the ligation in SnaBI-cut dephosphorylated vectors. The ligation mixtures were used to transform R1 competent cells. This was followed by the screening of clones using gene and vector-specific primers (gene Forward primer/ pET Reverse primer). For the expression of the protein, plasmid was used to transform BL21 (DE3) cells, and the protein induction was carried out with 0.2% arabinose for 4 h at 37 °C with streptomycin (50 µg/uL). RBD and human ACE-2 proteins expressed with a C-terminal hexa-Histidine tag were obtained in the inclusion bodies. For large scale purification, BL21 cells carrying RBD-pMTSA and ACE-2 pMTSA plasmid were grown in liquid culture overnight at 37 °C with streptomycin (50 µg/uL). The secondary culture was inoculated with 1% inoculum and grown till mid-log phase in the presence of streptomycin and induced with 0.2% arabinose for 4 h at 37 °C. The cell pellet was re-suspended in lysis buffer (50mM Tris pH8,10mM EDTA,100µg/ml lysozyme and 2mM PMSF) pH 8.0 and sonicated till a clear solution was obtained. The solution was subsequently centrifuged at 13,000 rpm for an hour to obtain a clear cell lysate and pellet. The pellet thus obtained, was solubilized overnight in solubilization buffer (10 mM Tris HCl, 150 mM NaCl, 8 M Urea pH 8). The solubilized pellet was again centrifuged at 13,000 rpm for 30 minutes to obtain a clear supernatant. The clear supernatant was allowed to bind Ni-NTA beads for 2h at RT, and the protein was subsequently purified by slowly increasing imidazole concentration in solubilization buffer. The eluted fractions were run on 15% SDS PAGE along with the Precision Plus Protein™ Dual Color Standards from Bio-Rad, and the fractions carrying the protein were pooled and refolded in refolding buffer (100mM Tris,20% glycerol,250mM L-arginine,1mM EDTA mixed and adjusted to pH8 and then added 1mM reduced glutathione (GSH) and 0.5mM glutathione disulfide) overnight at 4 °C. This was followed by dialysis in 10 mM Tris HCl pH 8 overnight at 4°C.

**Cloning, expression and protein purification of full-length Human Heat shock protein 70:** For the cloning of full length HSP70 in pMTSAra vector, inserts were prepared by digestion using SnaBI, followed by the ligation in SnaBI-cut dephosphorylated vectors. The ligation mixtures were used to transform R1 competent cells. This was followed by the screening of clones using gene and vector-specific primers (Full length Hsp70For/pET Rev). For the expression of the protein, HSP70-pMTSA plasmid was used to transform BL21 (DE3) cells, and the protein induction was carried out with 0.2% arabinose for 4 h at 37◦Full length Human HSP70 protein expressed with a C-terminal hexa-Histidine tag was obtained in the supernatant. The cell pellet was re-suspended in lysis buffer (50mM Tris pH8,10mM EDTA,100µg/ml lysozyme and 2mM PMSF) and sonicated till a clear solution was obtained. The solution was subsequently centrifuged at 13,000 rpm for an hour to obtain a clear cell lysate. The lysate was allowed to bind Ni-NTA beads overnight at 4◦C, and the protein was subsequently purified by slowly increasing imidazole concentration in 20mM

Tris HCl ,150Mm NaCl pH8. The eluted fractions were run on 15% SDS-PAGE, and the fractions having high concentrations of the protein were pooled and subsequently dialyzed against 10mM Tris. Buffer exchange was carried with PBS pH7.4. The purified protein was run on 15% SDS-PAGE.

**Surface Plasmon resonance**: Surface Plasmon resonance Real-time biomolecular interaction analysis with Surface Plasmon Resonance (SPR) was performed to determine and quantify the interaction between HSP70 and human ACE-2 using AutoLab Esprit SPR (Advanced Instrumentation Research Facility, Jawaharlal Nehru University, New Delhi, India), and the affinity constant, KD was established at Room Temperature (RT, 298 K). ACE-2 (30 μM in 10 mM Tris buffer, pH 8.0) was immobilized on a gold sensor chip (self-assembled monolayer of 11-Mercapto-Undecanoic Acid, MUA on gold surface) by covalent amine coupling. Full-length Hsp70 was injected individually at different concentrations (250 nM, 750nM, 1 μM, 2 μM, 4 μM and 6μM) over the ACE-2-immobilized chip surface, with the association and dissociation times of 300s and 200, respectively. Tris (10 mM, pH 8.0) was used for both immobilization and binding procedures.50 mM NaOH was used to regenerate the chip surface. The receptor binding domain of spike protein was injected individually at various concentrations (0.1μM,0.23μM,0.93μM,1.87μM and 7.5μM) over the ACE-2 immobilized chip surface, with the same association and dissociation time. This was followed by immobilization of Hsp70 (8 μM in 10 mM Tris buffer, pH 8.0) on the sensor chip surface, and the receptor binding domain of spike protein was injected at different concentrations (0.1μM,0.23μM,0.93μM,1.87μM and 7.5μM) over the Hsp70-immobilized chip surface. Finally, PES-Cl was injected at various concentrations (1 μM, 10 μM,15 μM, and 20 μM) onto the Hsp70-immobilized chip surface. Data were analyzed using Auto Lab ESPRIT Kinetic evaluation software.

**In silico docking analysis**

Docking of ACE-2 (PDB Id: 1R42) with human Hsp70 (PDB Id: 4PO2) was performed using HDOCK server. PyMol was used for visualization and analysis of protein structures.

**Supplementary Table 1.** Amino acid sequences of recombinant proteins (regions selected from spike protein, human ACE-2 receptor and HSP-70) expressed and purified for interaction studies (Surface plasmon resonance).

| Domain | Amino acid sequence | pI | GRAVY |
|---|---|---|---|
| Receptor binding domain SARS-CoV-2 (B.1.617.2) | MGIYQTSNFRVQPTESIVRFPNITNLCPFGEVFNATRFA SVYAWNRKRISNCVADYSVLYNSASFSTFKCYGVSPT KLNDLCFTNVYADSFVIRGDEVRQIAPGQTGKIADYNY KLPDDFT GCVIAWNSNNLDSKVGGNYNYRYRLFRKSN LKPFERDISTEIYQAGSKPCNGVEGFNCYFPLQSYGFQ PTNGVGYQPYRVVVLSFELLHAPATVCGPKKSTNLVK NKCVNFNFN | 9.08 | -0.326 |
| Human ACE-2 (Minimum region required to bind RBD) | MSTIEEQAKTFLDKFNHEAEDLFYQSSLASWNYNTNI TEENVQNMNNAGDKWSAFLKEQSTLAQMYPLQEIQ NLTVKLQLQALQQNGSSVLSEDKSKRLNTILNTMSTIY STGKV | 4.86 | -0.619 |
| Human HSP70 (Full length) | MAKAAAIGIDLGTTYSCVGVFQHGKVEIIANDQGNRTTP SYVAFTDTERLIGDAAKNQVALNPQNTVFDAKRLIGRK FGDPVVQSDMKHWPFQVINDGDKPKVQVSYKGETKA FYPEEISSMVLTKMKEIAEAYLGYPVTNAVITVPAYFND SQRQATKDAGVIAGLNVLRIINEPTAAAIAYGLDRTGKG ERNVLIFDLGGGTFDVSILTIDDGIFEVKATAGDTHLGG EDFDNRLVNHFVEEFKRKHKKDISQNKRAVRRLRTAC ERAKRTLSSSTQASLEIDSLFEGIDFYTSITRARFEELCS DLFRSTLEPVEKALRDAKLDKAQIHDLVLVGGSTRIPKV QKLLQDFFNGRDLNKSINPDEAVAYGAAVQAAILMGDK SENVQDLLLLDVAPLSLGLETAGGVMTALIKRNSTIPTK QTQIFTTYSDNQPGVLIQVYEGERAMTKDNNLLGRFEL SGIPPAPRGVPQIEVTFDIDANGILNVTATDKSTGKANKI TITNDKGRLSKEEIERMVQEAEKYKAEDEVQRERVSAK NALESYAFNMKSAVEDEGLKGKISEADKKKVLDKCQEV ISWLDANTLAEKDEFEHKRKELEQVCNPIISGLYQGAG GPGPGGFGAQGPKGGSGSGPTIEEVD | 5.61 | -0.417 |

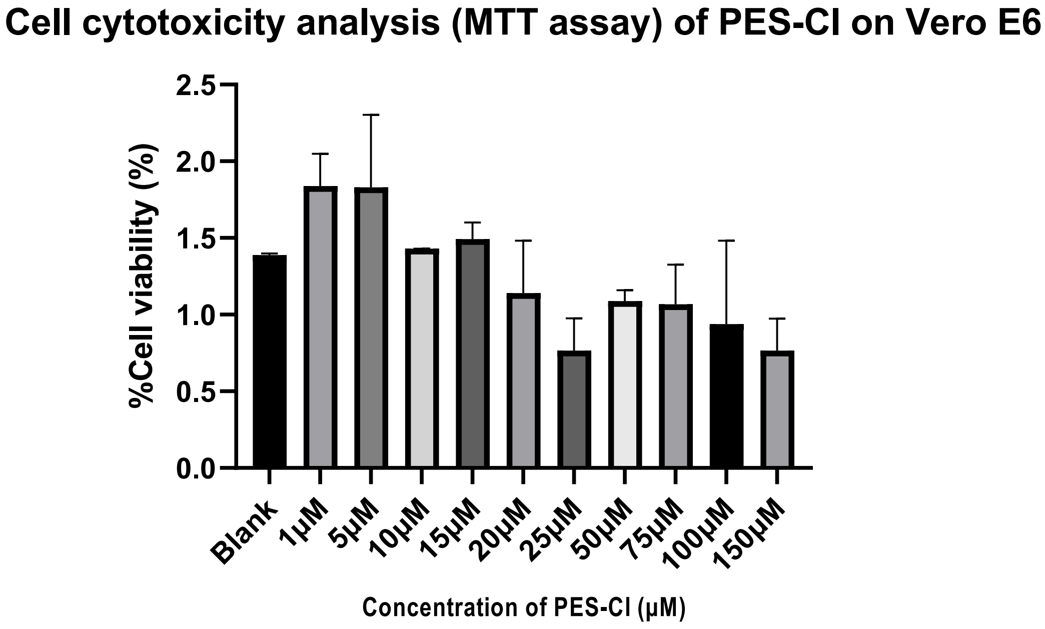


Supplementary Figure 1. Cell cytotoxicity analysis (MTT assay) of PES-Cl on Vero E6 cells after 48 hours of treatment at indicated concentrations.

**kD= 100 µM**

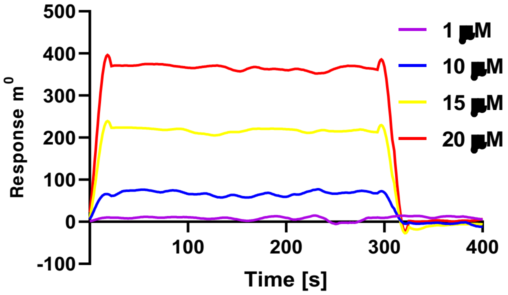


Supplementary Figure 2. Quantification of the interaction strength. SPR-based interaction analysis upon titrating the immobilized Hsp70 with PES-Cl. Kd value is indicated above the sensograms.

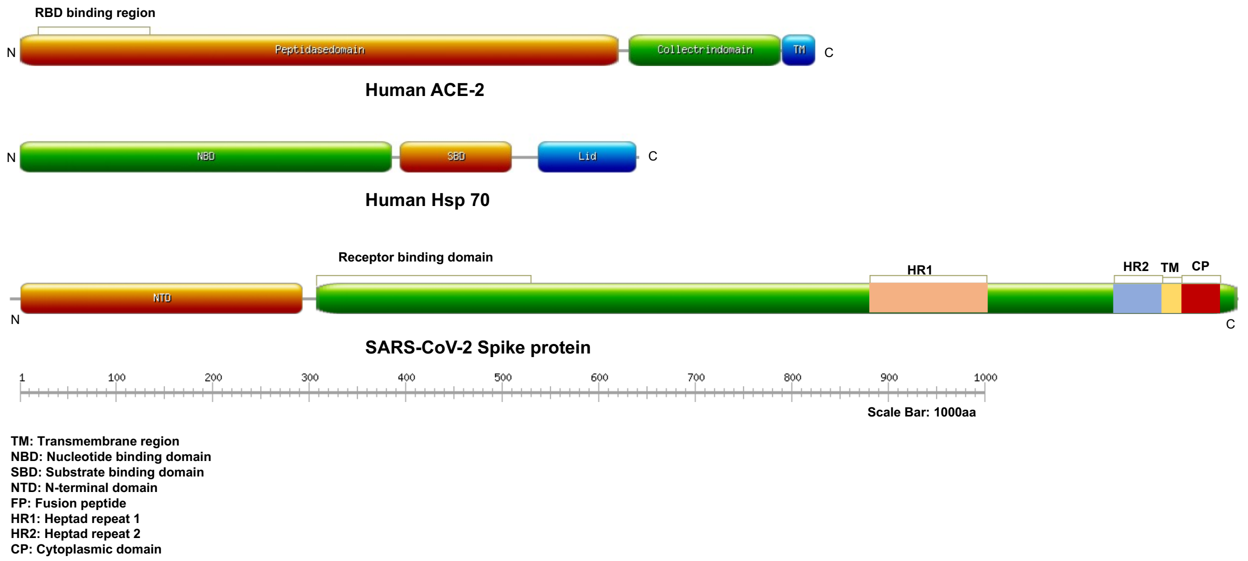


Supplementary Figure 3. Domain architecture of Human ACE-2, Spike protein of SARS-CoV2 and human HSP70. The boxed line above the schemes (RBD binding region in ACE-2 and Receptor Binding Domain in Spike protein), indicate the region which is recombinantly expressed and purified for this study. For Hsp70, full length protein is recombinantly expressed and purified.

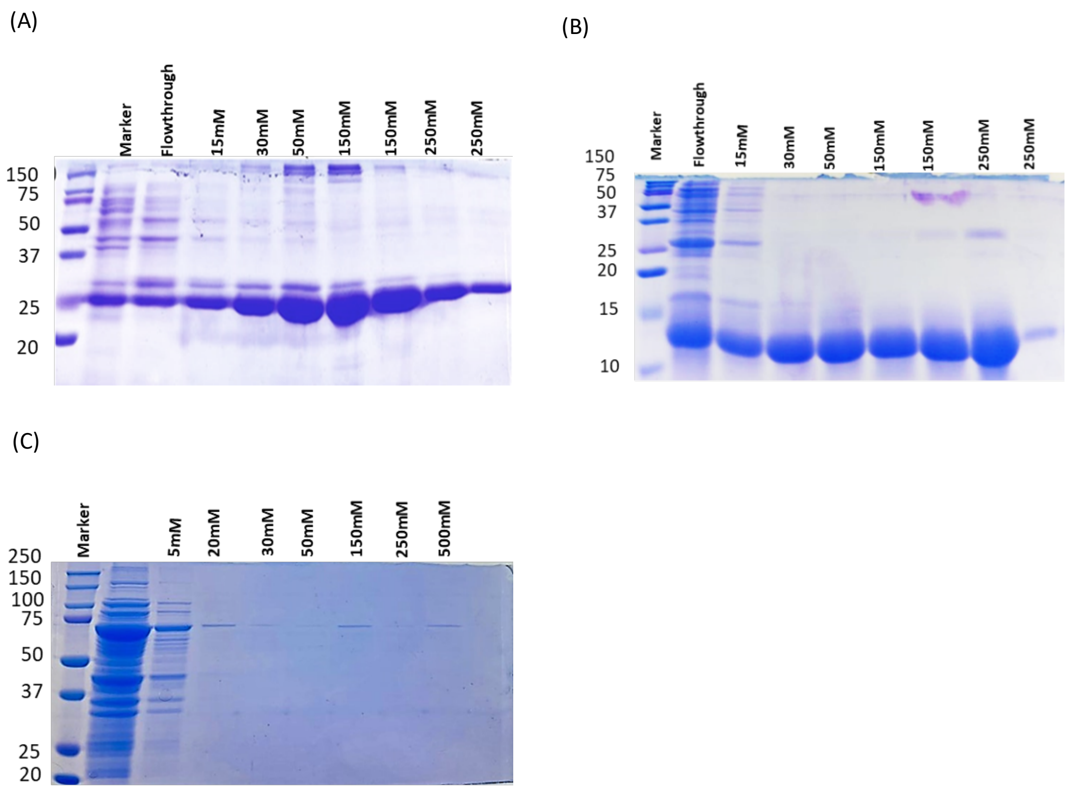


Supplementary Figure 4. (A) ACE-2 purification from inclusion bodies by Ni-NTA affinity chromatography at different concentrations of Imidazole. (B) RBD purification from inclusion bodies by Ni-NTA affinity chromatography at different concentrations of Imidazole. (C) Full-Length HSP70 purification from supernatant by Ni-NTA affinity chromatography at different concentrations of Imidazole.

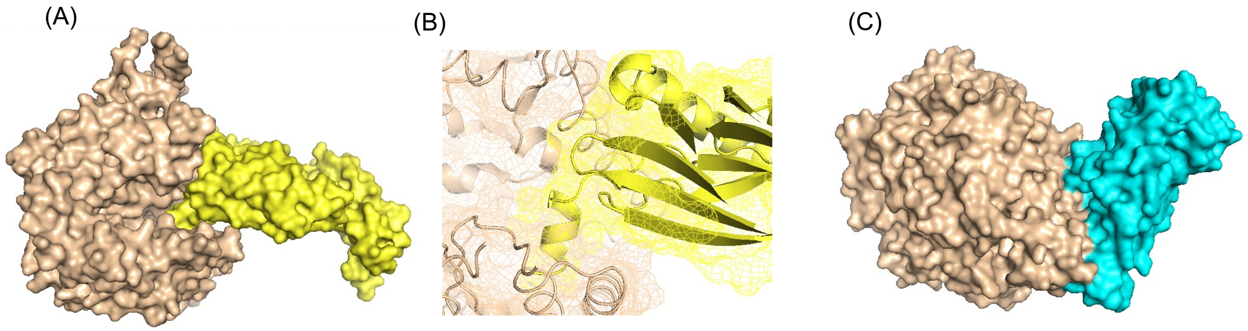


Supplementary Figure 5: (A) Surface representation of docked complex of ACE2 with human Hsp70. ACE2 and human Hsp70 are shown in wheat and yellow color respectively. (B) Zoom image representing the interaction interface of ACE2-Hsp70 docked complex. (C) Surface representation of crystal structure of chimeric omicron RBD (strain BA.1; represented in wheat color) complexed with human ACE2 (denoted by cyan color).

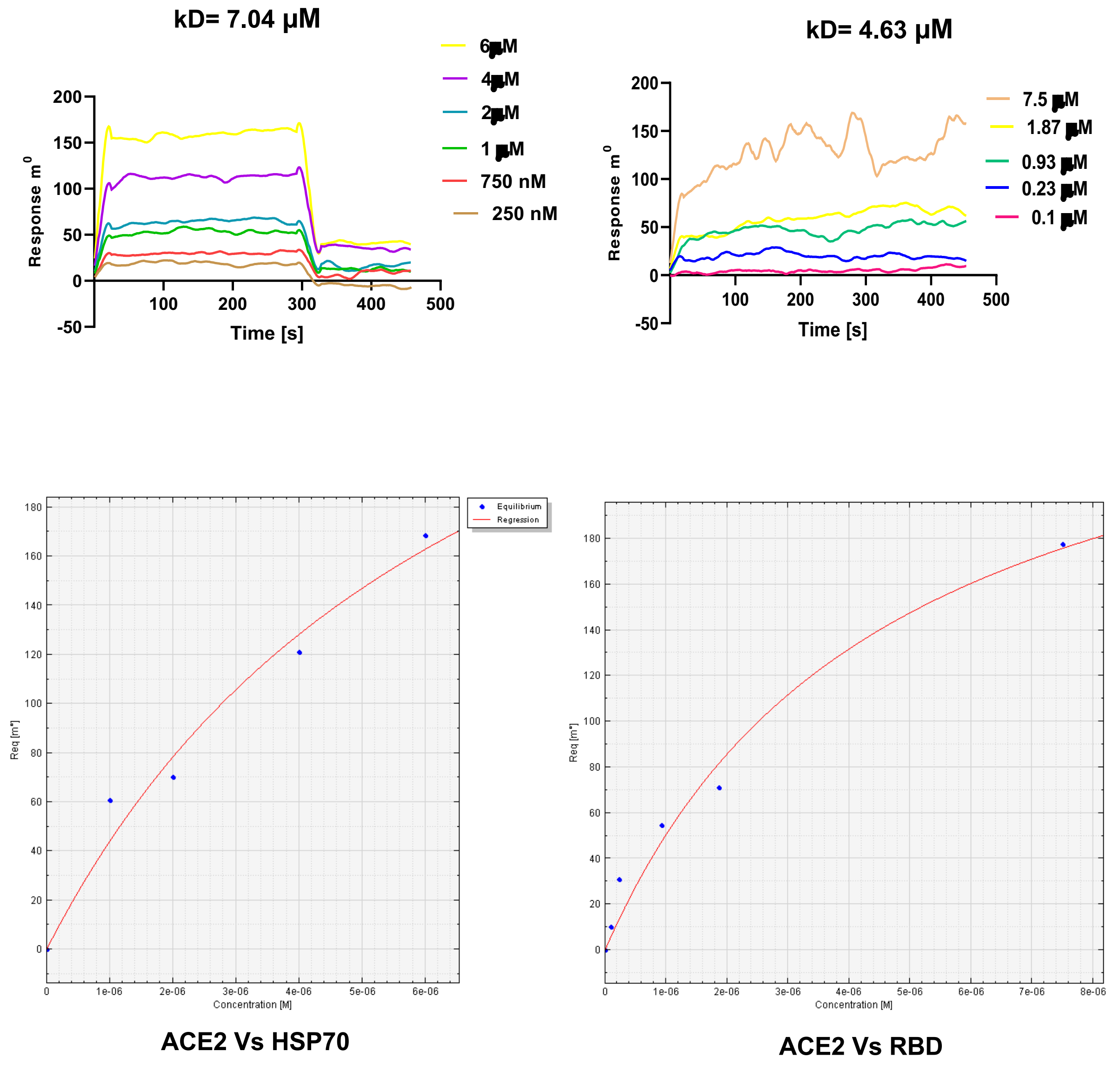


Supplementary Figure 6. Quantification of the interaction strength. SPR-based interaction analysis upon titrating the immobilized ACE-2 with HSP70 and RBD domain of spike protein. The Kd values are indicated above the sensograms.
